## Supplemental data for "RPRM as a potential preventive and therapeutic target for radiation-induced brain injury via multiple mechanisms"

*To whom correspondence should be addressed: H. Yang

**Running title:** RPRM as a target for IR-induced brain injury

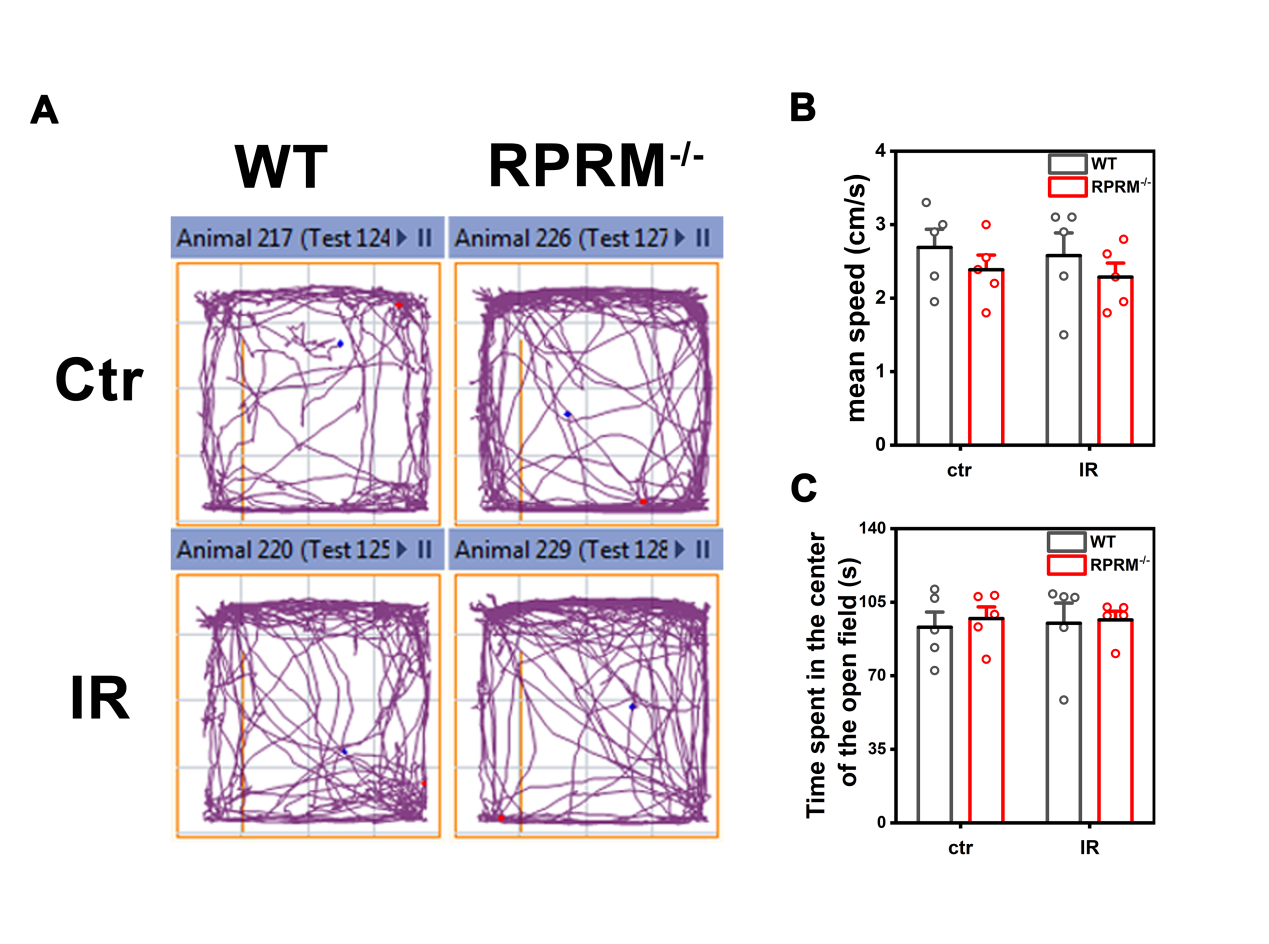

**Fig. S1** The open field test on 6–8-week-old WT and RPRM KO male mice after exposed to 10 Gy WBI. **A** Typical exploration tracks of the four groups of mice in the open field test. **B** Mean speed of the four groups of mice in the open field test. **C** Time the four groups of mice spent in the center of the open field. n=5. Data were presented as mean ± SEM. Data were analyzed using two-way ANOVA with Tukey’s correction.

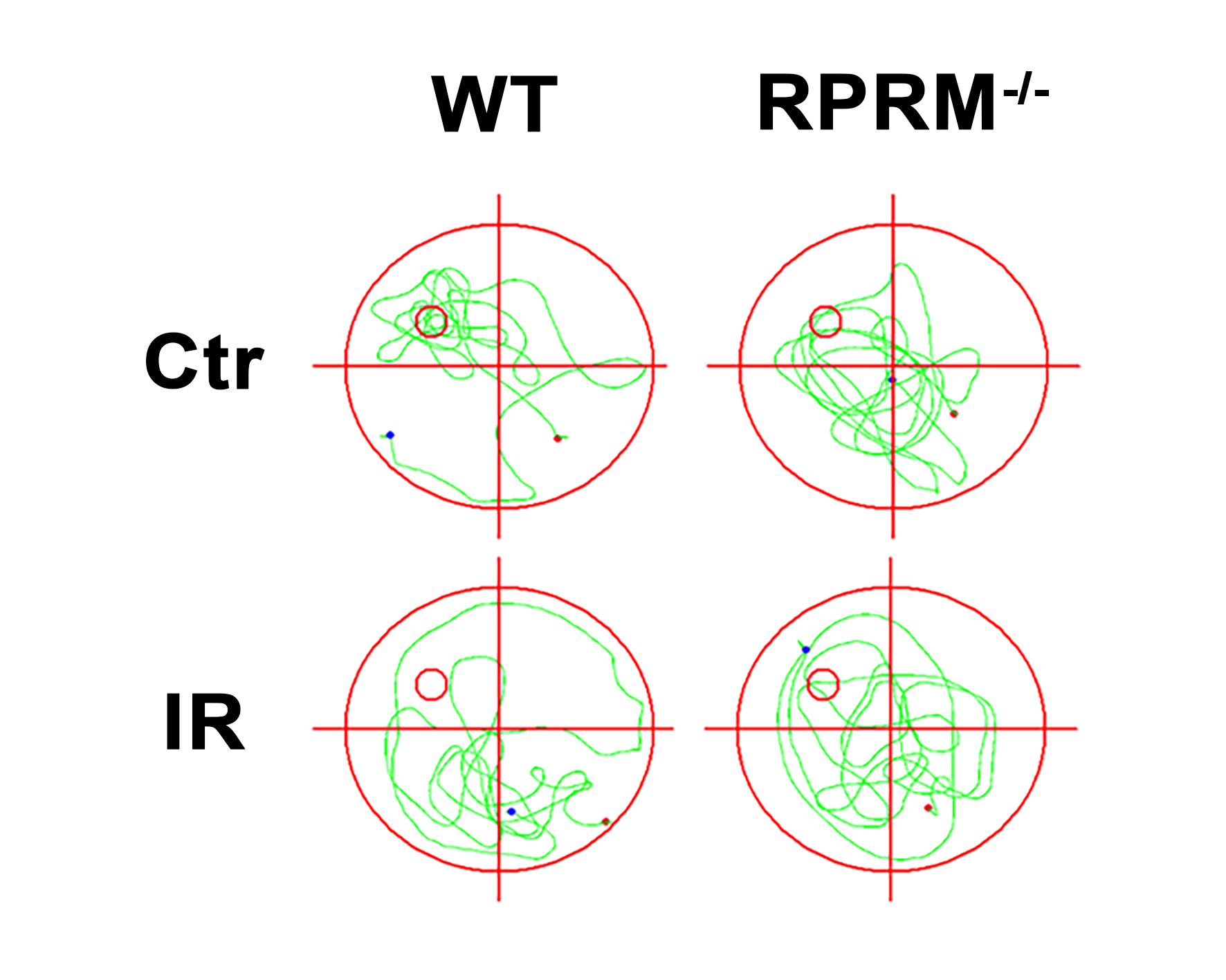

**Fig. S2** Representative exploration tracks of 6–8-week-old WT and RPRM KO mice in the spatial probe test of MWM 2 months after exposed to 10 Gy WBI.

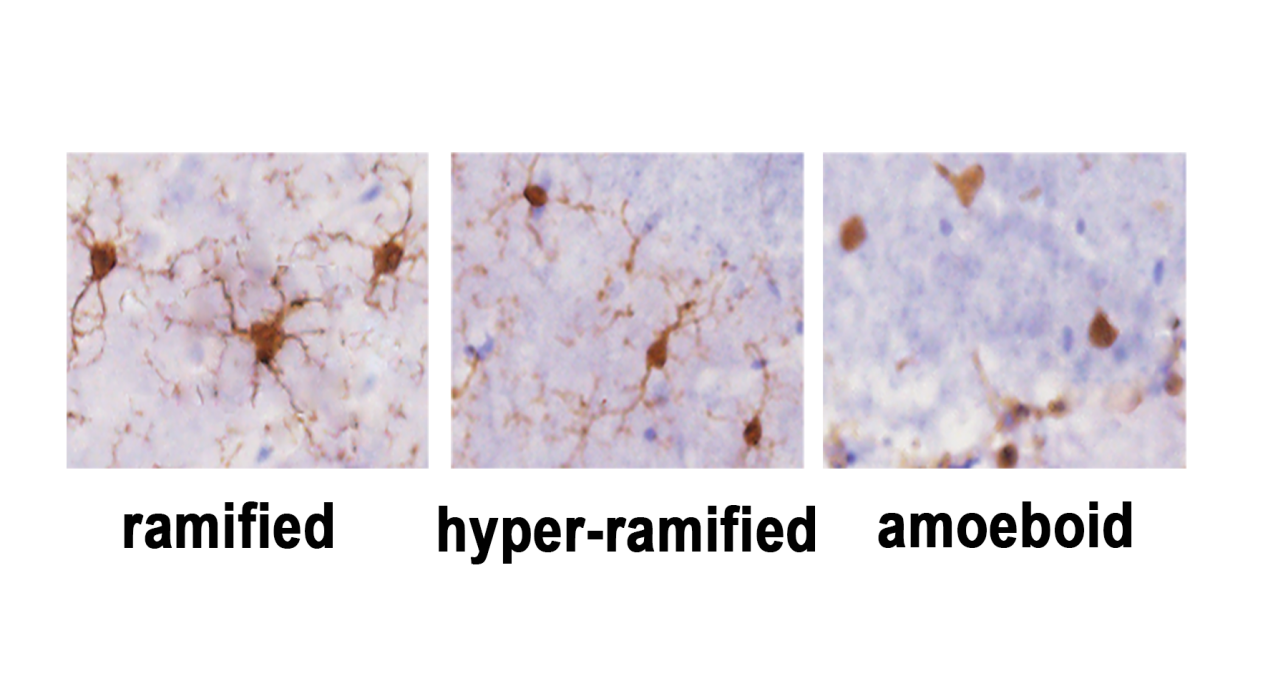

**Fig. S3** Representative morphology of microglia in different status including ramified (non-activated), hyper-ramified (reactive) and unramified (activated) in mouse hippocampus.
